# Organotypic slice culture model demonstrates interneuronal spreading of alpha-synuclein aggregates

**DOI:** 10.1101/681064

**Authors:** Sara Elfarrash, Nanna Møller Jensen, Nelson Ferreira, Cristine Betzer, Jervis Vermal Thevathasan, Robin Diekmann, Mohamed Adel, Nisreen Mansour Omar, Mohamed Z. Boraie, Sabry Gad, Jonas Ries, Deniz Kirik, Sadegh Nabavi, Poul Henning Jensen

## Abstract

Here we describe the use of an organotypic hippocampal slice model for studying α-synuclein aggregation and inter-neuronal spreading initiated by injection of preformed α-synuclein filaments (PFFs). PFF injection at dentate gyrus templates the endogenous α-synuclein to form aggregates in axons and cell bodies that spread to CA3 and CA1 regions. Aggregates were insoluble and phosphorylated at serine 129, recapitulating Lewy pathology features found in Parkinson’s disease and other synucleinopathies. The spreading of the aggregates were favoring the anterograde direction in the slice model. The model allowed development of slices expressing only serine-129 phosphorylation-deficient human α-synuclein (S129G) using adeno-associated viral (AAV) vector in α-synuclein knockout slices. Processes of aggregation and spreading of α-synuclein were thereby shown to be independent of phosphorylation at serine 129. We provide methods and highlight crucial steps for PFF microinjection and characterization of aggregate formation and spreading. Slices derived from genetically engineered mice or manipulated by using viral vectors allow testing of hypotheses on mechanisms involved in formation of α-synuclein aggregates and their prion-like spreading.

## 1. Introduction

Parkinson’s disease (PD) is characterized by appearance of abnormal proteinaceous inclusions, Lewy bodies, whose development progress through the nervous system. Lewy bodies have been hypothesized to originate in the gut and olfactory bulb and spread from there through vulnerable neuronal populations in the brain (Braak et al., 2003),(Brundin & Melki, 2017). Alpha-synuclein (α-syn) is a presynaptic protein of unknown function that was reported to be the main component of the Lewy body (Spillantini et al., 1997). The progressive spreading of α-syn containing Lewy pathology in different brain regions of PD patient’s, has suggested an intercellular transfer of seeding-competent α-syn aggregates from an affected neurons to a healthy one, that upon uptake, templates aggregation of native α-syn into toxic and seeding-competent aggregates that can perpetuate the process through the nervous system (Braak et al., 2003),(Jucker & Walker, 2018),(Kara, Marks, & Aguzzi, 2018).

This hypothesis is supported by several *in vivo* experiments where injection of seeds, consisting of preformed α-syn fibrils or α-syn aggregate-containing brain extracts had induced spreading of α-syn pathology (Jucker & Walker, 2018),(Rey et al., 2016),(Mougenot et al., 2012),(Luk et al., 2012). It is also supported by cell-based models allowing mechanistic studies of uptake, seeded aggregation, intercellular spreading and drug screening. (Karpowicz et al., 2017), (Volpicelli-Daley et al., 2011),(Tran et al., 2014).

However, brain complexity in *in vivo* models and the simplicity of cell-based models hamper the study of mechanisms operating in the brain and invite such basic questions as: Is there a preferred, e.g. trans-synaptic, mode of spreading in relation to anterograde or retrograde axonal transport processes, or is it less specific where seeds are released from the somato-dendritic compartments of the affected neurons?

Here, we introduce a novel brain tissue model that allows inter-neuronal spreading of α-syn aggregate pathology to be studied on a faster time scale than in *in vivo* models and with the ease of manipulation observed in cell-based models. The model combines organotypic mouse hippocampal cultures with region-specific injection of preformed α-syn fibrils (PFFs). Using hippocampal slices from wild type, α-syn-transgenic and α-syn-knockout (KO) pups in combination with viral vector-based expression of wild type and non-phosphorylatable S129G α-syn, we demonstrate that the preferred mode of spreading is an anterograde trans-synaptic transfer requiring no Ser129 phosphorylation, albeit this post-translational modification represents a marker of Lewy bodies.

The swiftness of the templated α-syn aggregates development and spreading in the model combined with its flexibility with respect to using tissue from genetically modified mouse strains and viral vector technology opens up novel strategies for investigating molecular mechanisms operating in the spreading of α-syn pathology in the brain. Adopting such live-cell approaches and using *in vivo* imaging and electrophysiology, the model may be used for studying the impact of templated α-syn aggregation on neuronal functionality and connectivity.

## 2. Materials and Methods

### 2.1. Preparation of organotypic hippocampal slice cultures

Organotypic hippocampal slices were made from 5-7-day-old post-natal pups from C57BL/6 (wild type), SNCA^−/-^ α-syn KO (C57BL/6N-Sncatm1Mjff/J from Jackson lab), or mThy-1-human α-syn-expressing mice (Tg(Thy1-SNCA)61Ema) (Rockenstein et al., 2002) according to Stoppini et al. 1991(Stoppini, Buchs, & Muller, 1991). Low Na cerebrospinal fluid (CSF) was carbogenated on ice until the color changed to orange and some ice slashes were formed (1 mM CaCl_2_, 10 mM D-glucose, 4 mM KCl, 5 mM MgCl_2_, 26 mM NaHCO_3_, 234 mM sucrose and 0.1% phenol red solution). 15 ml of low Na CSF was prepared for each brain in a 50ml Falcon tube. After decapitation of the pups, the extracted brain was moved gently and kept in low Na CSF for 1 minute, before the brain and the low Na CSF were poured gently into a petri dish for hippocampus dissection under microscopic guidance (SZX-ZB7 Stereomicroscope, Olympus x1 objective). Slices of 400 µm thickness were made using a tissue chopper (Stoelting, #51425) before being moved to a culture medium (MEM Eagle medium 78.8% (Gibco #11095), 20% heat-inactivated horse serum (Gibco, #16050-122), 1 mM L-glutamine,1 mM CaCl_2_, 2 mM MgSO_4_, 170 nM insulin, 0.0012% ascorbic acid, 12.9 mM D-glucose, 5.2 mM NaHCO_3_, 300 mM Hepes (Sigma #H3375), pH=7.28, osmolality adjusted to 317-322) pre-heated to 37°. Only slices with intact DG and CA regions were selected under the microscope and moved to air-fluid interface-style Millicell culture inserts, 30 mm diameter, 0.4 µm (Millipore) in 6-well culture plates (ThermoFisher Scientific) with 800 µL of sterile medium added below the insert (Stoppini et al., 1991). The full medium was changed trice weekly. All steps of the procedure after decapitation and brain extraction were performed in a laminar flow tissue culture hood using sterile equipment and aseptic technique.

### 2.2 Preparation of S129A α-syn pre-formed fibrils (PFFs)

Recombinant human α-syn with residue serine129 to an alanine was expressed in E. coli and purified as described in wild type α-syn (Huang, Ren, Zhou, & Wang, 2005). To make pre-formed S129A-α-syn PFFs, monomeric S129A α-syn (5 mg/ml) in Eppendorf tubes was incubated in sterile phosphate-buffered saline (PBS), pH 7.4 (Gibco), for 48 hours at 37°C on a Thermomixer (Eppendorf) at 1050 rpm. To validate sufficient aggregation, 50 µl of the incubate was centrifuged at 25,000 × *g* for 20 min and the supernatant was isolated. PBS, 50 µl, was added on the pellet and 50 µl 2 x SDS-loading buffer (20 mM Tris, pH 6.8, 2 mM EDTA, 80 mM DTT, 2% SDS, 20% sucrose) was added to the pellets and supernatants, which were heated to 96°C for 15 min. Equal volumes of supernatants and pellets were subjected to sodium dodecyl sulfate polyacrylamide gel electrophoresis (SDS-PAGE) analysis on 8-16% Bis-Tris gels (Genscript) that subsequently were stained with Coomassie blue R-250 (Supplementary fig. 1,a,i). Approximately 75% of the protein was routinely recovered in the insoluble pellet fraction. To ascertain the development of amyloid-type aggregates in the insoluble fractions, K114 fluorometry was conducted (Crystal et al., 2003). Equal volumes of aggregated and monomer α-syn S129A were mixed with the (trans,trans)-1-bromo-2,5-bis-(4-hydroxy)styrylbenzene (K114) (50 µM) in 100 mM glycine, pH 8.5, after which fluorescence (λ_ex_=380 nm, λ_em_=550 nm, cutoff = 530 nm) was measured with a Perkin Elmer, EnSpire 2300 Multilabel Reader (Supplementary fig. 1,a,ii).

When proper aggregation had been assured, the insoluble S129A α-syn was isolated by centrifugation in Eppendorf tubes. After discarding the supernatant, the pellet containing the PFFs was diluted to 2 mg/ml in sterile PBS, pH 7.4 (Gibco) and subjected to ultrasound breakage for 20 min using a sonicator (Branson 250, settings: 30% Duty Cycle, Output Control 3) equipped with a water jacket cooling system to avoid sample heating during sonication. The size distribution profile of the PFFs was measured using dynamic light scattering (DLS) using a Wyatt DynaPro NanoStar instrument at 25°C. Data were processed using the Dynamics 7.5.0.17 software package with the solvent (PBS) background signal subtracted from each sample. The PFF sample showed a homogeneous, monodispersed population with a hydrodynamic radius of 44 nm (Supplementary fig. 1,a,iii). The PFFs were dispensed into sterile Eppendorf tubes in (75 µl) aliquots of concentration, 2mg/ml, snap frozen and stored at −80 ° C. A sample was used to validate the protein concentration using BCA analysis.

### 2.3. Microinjection of organotypic slices

To facilitate injection, each insert was moved from the 6-well plate to a sterile cell culture dish, 35 x 10 mm (SARSTEDT #83.3900.500), before injection. Slices were microinjected in the DG after 7 days in culture, except where otherwise mentioned. Light microscopy was used to identify DG by a characteristic horseshoe arrangement of the nuclei of granule cells.

Immediately before injection, the aliquot of S129A PFFs was thawed at room temperature (RT) and sonicated for 30 seconds using the above settings (Branson, Sonifier 250). After sonication, PFFs were kept at RT during the injection process. Microinjection pipette (item #1B200F-4 (with Filament), WPI) was pulled using a micropipette puller (P-1000, Sutter Instrument). For microinjection, a Pulse Pal v2 (#1102) was used (settings: phase 1 voltage 5V, phase 1 duration 0.01 seconds, pulse interval 0.5 seconds).

Injections were performed in a laminar flow hood equipped with a microscope to ensure aseptic conditions. The pipette was loaded using Eppendorf microloader pipette tips (ThermoFisher); the pipette was inserted into the holder, and the tip was cut with a fine scissor under visual guidance. Pressure pulse was applied to test whether the PFF suspension was expelled from the tip. Once the tip was open, a small droplet should emerge. Correct injection was confirmed by short lifting of the surface of the tissue at the injection site. After injection, the needle was left in place for 20 seconds and then slowly removed. The volume injected in the slices was estimated by counting the number of shots with the adjusted Pulse Pal set up (10-12 shots/slice, 10 nl/shot, 0.1 μg/slice). The final volume was injected at the DG at two to three injection sites depending on the slice thickness at the site of needle insertion. It was important to pay attention to tissue architecture under microscopic guidance during the injection procedure to avoid injection of a large volume at a single site which would cause rupture of the tissue and release of the PFFs to the surface of OHCS. The session lasted approximately 6 to 8 minutes for an insert holding four slices. The final volume injected in each slice was about 0.1µl of either S129A-PFF (1 mg/mL), monomeric α-syn (1 mg/mL), or PBS. After injecting all slices on a culture insert, the medium was replaced with a fresh pre-heated medium.

### 2.4. Adeno-associated viral (AAV) mediated expression of wt and S129G α-syn in slices from α-syn KO mice

Pseudotyped rAAV2/6 WT human α-syn and rAAV2/6 S129G human α-syn vectors were produced by using a co-transfection method with rAAV transfer plasmid containing the gene of interest placed between two AAV2-inverted terminal repeats and a helper plasmid (pDP6) coding for necessary elements for production and packaging of the capsid particles. The gene of interest was driven under the control of a human synapsin-1 promoter. Vectors were purified by iodixanol step gradients and ion exchange chromatography, as described in detail elsewhere (Zolotukhin et al., 2002). The final titers used in the experiments were 3.5 x 10E13 genome copy (gc)/ml. All titers were determined by quantitative PCR using TaqMan probes targeting the ITR sequences. After 3 days in culture, AAV-human-α-syn and AAV-human-S129G-α-syn vectors were injected in slices made from P5 α-syn KO pups. The slices were either injected with the virus at the three regions DG, CA3, and CA1 or only at DG and CA1. This procedure expresses α-syn either in the three synaptically connected regions or leaves the interconnecting CA3 region without α-syn expression. Three days after AAV injection, α-syn S129A PFFs were injected at DG as demonstrated in Fig. 3c,d.

### 2.5. Immunohistochemistry

Organotypic slices were fixed using 4% PFA in phosphate buffer (PB) (2.8 mM NaH_2_PO_4_H_2_O, 7.2 mM Na_2_HPO_4_.2H_2_O, 123 mM NaCL, pH adjusted to 7.2) and processed for immunohistochemistry according to Gogolla et al., with slight modifications (Gogolla, Galimberti, DePaola, & Caroni, 2006). Briefly, after fixation, slices were permeabilized using 0.5% Triton X-100 for 6 hours at RT or overnight at 4°C with slight shaking. Slices were then incubated with blocking buffer (10% bovine serum albumin (BSA)/PB) for 3 hours at RT. The primary antibody was prepared in 5% BSA/PB and incubated with slices overnight at 4°C during gentle shaking. Antibodies used were α-syn aggregate-specific antibody MJF14 (rabbit monoclonal MJFR-14-6-4-2. Abcam,1:25,000), the two phospho-serine129-specific antibodies, 11A5(Anderson et al., 2006) (mouse monoclonal 11A5 kindly provided by Imago Pharmaceuticals, 1:25,000) and D1R1R (rabbit mAb #20706S, Cell signaling. 1:1000), NF-L (mouse mAb #2835, Cell signaling. 1:500) NeuN (mouse mAb clone A60, Millipore. 1:200), and anti-alpha-synuclein antibody MJFR1 (rabbit monoclonal ab138501, Abcam, 1:5000). The slices were washed trice using TBS (NaCl 150 mM, Tris 20 mM) with 0.3% Triton X-100 washing buffer during gentle shaking for 30 minutes/wash. After the final wash, the slices were incubated with the appropriate Alexa Fluor dye (488 and 568) labeled secondary antibody (Invitrogen 1:2000) and 4’,6 diamidino-2 phenylindole (DAPI) (TH.GEYER, 5 mg/mL, 1:1000) in 5% BSA/PB for 3 hours at RT during gentle agitation shielded from light. The slices were washed trice as above and mounted using DAKO mounting medium (DAKO, D3023). The edges of the coverslip were sealed using nail polish.

The staining could be done either by i) directly adding reagents to the inserts (1ml about and 1ml below the insert) or ii) excising the slices from the inserts with their culture membrane below with a scalpel and further incubating them with antibodies in 96-well plates to save reagent. For the washing steps, 24-well plates were used to ensure proper rinsing.

### 2.6. Preparation of the organotypic hippocampal slice culture for super-resolution dSTORM imaging

For cryosection, 4% PFA-fixed slices were kept in 30% sucrose at 4°C overnight for cryoprotection. The slices were cut out of the membrane insert with a piece of membrane below. Holding the slice carefully from the membrane, the slices were mounted on the cryostate stage using Tissue-Tek® O.C.T. Compound (Sakura), where the slice surface faces downwards. After the OCT solidified, the membrane was carefully removed and an extra amount of OCT was added on top of the slice and cut on cryostate (Leica) at a thickness of 10 µm at −20 °C. The sections were collected on Superfrost Plus Adhesion Microscope Slides (ThermoFisher). Immunostaining was processed on the slides as previously described for the whole mount using an antibody against pS129. During antibody incubation, slides were kept in a humidity chamber with a hydrophobic pen barrier erected around the tissue to prevent it from drying out during incubation. After the final washing step, the slides were mounted using glycerol gelatin aqueous slide mounting medium (Sigma).

#### Tissue preparation for super-resolution dSTORM imaging

Stained tissue slices were gelatin-embedded and sandwiched between a rectangular coverslip (50 X 24 mm) and microscope slide (75 X 25 mm). To facilitate dSTORM imaging, conventional dSTORM imaging buffer (50 mM Tris/HCl pH 8, 10 mM NaCl, 10% (w/v) D-Glucose, 500 µg/mL glucose oxidase, 40 µg/mL glucose catalase and 35 mM MEA in H_2_O) was added to the embedded slice. The rectangular coverslip was carefully removed by immersing the microscope slide into a beaker of PBS prewarmed to 60 °C. Imaging buffer (100 uL) was added to the gelatin layer, and a new rectangular coverslip was then placed over the gelatin layer. The coverslip edges were then sealed using a 2-component silicon glue. Once the glue was set, the coverslip and the microscope slide sandwich were mounted on a microscope stage for imaging.

#### Microscope setup for dSTORM acquisition

dSTORM image acquisition was performed on a custom-built inverted microscope. Laser light at 405 nm and 640 nm wavelengths emitted from a laser box equipped with a single-mode fiber (iChrome MLE, Toptica) was collimated using an achromatic lens (f = 30 mm, Thorlabs), relayed by a 4f microscope of two lenses (each f = 250 mm, Thorlabs), spatially filtered for background reduction by an iris conjugate to the image plane (SM1D12D, Thorlabs), and focused onto the back-focal plane of a water-immersion objective lens (60x / NA1.2, Olympus). Mounting the fiber on a one-axis stage (SLC2445me-4, Smaract) allowed for image acquisition in HILO (highly inclined and laminated optical sheet) mode (Tokunaga, Imamoto, & Sakata-Sogawa, 2008). Fluorescence excitation and emission were separated using a multi-line dichroic beam splitter (zt405/488/561/640rpc, Chroma). Fluorescence emission was additionally filtered by a band pass filter (676/37 BrightLine HC, AHF) and directly focused onto an sCMOS camera (Orca Flash 4.0, Hamamatsu) via a tube lens (f = 180 mm, MVX-TLU, Olympus) resulting in projected pixel widths of 108 nm. Raw dSTORM data were recorded at a frame rate of 40 Hz and 25 msec exposure time. The lasers were triggered using an FPA (Mojo, Embedded Micro) controlled by a custom-written Micro Manager 1.4.22 plugin (Mund et al., 2018). This kept the number of activated emitters per frame approximately constant to a preset value via modulation of the pulse length of the 405 nm laser. The intensity of the fluorescence excitation laser at 640 nm was about 10 kW/cm^2.

### 2.6. Immunoblotting analysis of brain slice extracts

Slices were collected by cutting the tissue with a piece of its supporting culture insert membrane to ensure that all tissue was collected. For each time point, eight slices were collected. After 14 days in culture (7dpi of the PFFs), around 30 μg of protein was extracted per slice.

After washing twice in Hank’s buffer (5.37 mM KCL, 0.44 mM KH_4_PO_4_, 0.44 mM Na_2_HPO_4_, 136.9 mM NaCL), the tissue was homogenized with a tissue homogenizer (VWR #4320202) in ice-cold radioimmunoprecipitation assay (RIPA) buffer (50 mM Tris (pH 7.4), 150 mM NaCl, 1% Triton X-100, 2 mM EDTA, 0.5% sodium deoxycholate, 0.1% SDS) supplemented with protease inhibitor cocktail (Complete, Roche) and phosphatase inhibitors (25 mM B-glycerolphosphate, 5 mM NaF, 1 mM Na_3_VO_4_, 10 mM Na-pyrophospate). After homogenization, samples in Eppendorf tubes were sonicated (Branson, Sonifier 250, settings: 30% Duty Cycle, Output Control 3, 40 shots) before centrifugation at 25,000 x g for 25 min at 4°C. Supernatants were collected as the RIPA-soluble fraction. The pellets were resuspended in RIPA buffer, which was washed twice by centrifugation to remove remaining soluble material from the insoluble pellets. The pellets were dissolved in SDS-urea buffer (4% SDS, 50 mM Tris, 7 M urea, 40% glycerol and bromophenol blue, 2.5 mM DTE) overnight at RT as the RIPA-insoluble aggregate fraction. Protein concentration was measured using the bicinchoninic acid assay (BCA) kit (Sigma).

The RIPA-soluble fraction lysates were supplemented with an SDS-PAGE loading buffer (50 mM Tris pH 6.8, 4% SDS, 2.5 mM DTE, 40% glycerol, bromophenol blue), whereas the RIPA-insoluble fraction was ready for loading on the gel. Samples were heated to 95°C for 5 min and resolved on 8-16% Bis-Tris gels (Genscript) before being transferred to PVDF membranes using the Iblot2 Dry blotting system (ThermoFishcer). Membranes were fixed in 4% PFA for 30 minutes and blots probed for α-syn were boiled in PB for 10 minutes to improve immunodetection of α-syn (Lee & Kamitani, 2011). Membranes were blocked for 1 hour at RT in blocking buffer (5% skimmed milk powder, 20 mM Tris base, 150 mM NaCl, 0.05% Tween 20-containing phosphatase inhibitors) supplemented with 0.02 % NaN_3_. Primary and secondary antibodies were diluted in blocking buffer. Incubation with the primary antibody was done overnight at 4°C and with secondary antibodies (DAKO #P0217, #P0260) for 1.5 hours at RT with washing in TBS-Tween trice for 15 minutes each after each incubation. Bound antibodies were visualized using enhanced chemiluminescence (ECL) in a Fuji LAS-3000 Intelligent Dark Box (Fujifilm, Japan). To reprobe filters, they were stripped for bound antibodies to the PVDF membranes using Restore Western Blot Stripping Buffer (Thermoscientific, #21059) according to the manufacturer’s recommendation. Membranes were then processed with blocking and antibody detection as mentioned above. Antibodies used were the following: rabbit polyclonal anti-α-syn (ASY-1) 1:1000 (Jensen et al., 2000), rabbit monoclonal anti-α-syn antibody (MJFR1,ab138501, 1:1000), mouse monoclonal pS129-α-syn (11A5 kindly provided by Imago Pharmaceuticals, 1:2000), mAb anti-β-Tubulin III (TUJ1, T8578, Sigma, 1:5000), mAb mouse-specific α-syn (D37A6) (XP Rabbit #4179, CS 1:1000), mAb anti-α-Syn Syn-1 (BD Transduction Laboratories, 1:1000). PageRuler pre-stained protein ladder 10-180 kDa (ThermoFischer, #26616) was used as the molecular size marker.

### 2.7. Quantification

Quantification of Western blots was done using ImageJ (National Institutes of Health) after first assuring that the bands were not saturated. For quantification of immunostainings, four pictures covering the whole organotypic slice were taken using x10 objective, and a threshold was set where only the aggregate-specific signals were visible. The same threshold was applied to all images. For analysis, the mean fluorescence intensity (MFI) of the selected aggregate signals was quantified using ImageJ (National Institutes of Health) software. Signals were normalized to the total surface area of each slice detected using the DAPI staining.

### 2.8. Statistical data and analysis

Statistical analysis was performed using unpaired Student’s T-test for comparison of two groups. Data are presented as means +/-standard deviation (SD) *p<0.05, **p<0.01, ***p<0.001. For dSTORM data analysis, all data analysis and image reconstructions were performed with custom software written in MATLAB which is available as open source (github.com/jries/SMAP). Single molecule events were localized using a Gaussian fitter. Reconstructed images were rendered after filtering the localization table based on localization precision and point spread function width.

## 3. Results

### Progressive accumulation of mouse α-syn in organotypic hippocampal slice cultures and efficient C-terminal truncation of injected preformed α-syn fibrils

To study α-syn aggregate pathology and its spreading in brain tissue, we developed a new *ex vivo* model based on the organotypic mouse hippocampal slice culture method. The slices were prepared from 5 to 7-days-old post-natal pups of wild type BL6 mice, cultivated on a membrane by the air-fluid interphase method (Stoppini et al., 1991) (Fig. 1, b). This preparation has been extensively used for electrophysiological studies because it exhibits well-characterized synaptic connectivity between granule neurons in DG and CA3 pyramidal neurons that subsequently form synapses on CA1 pyramidal neurons (Dunwiddie & Lynch, 1978) (Fig. 1, a).

**Fig. 1:**
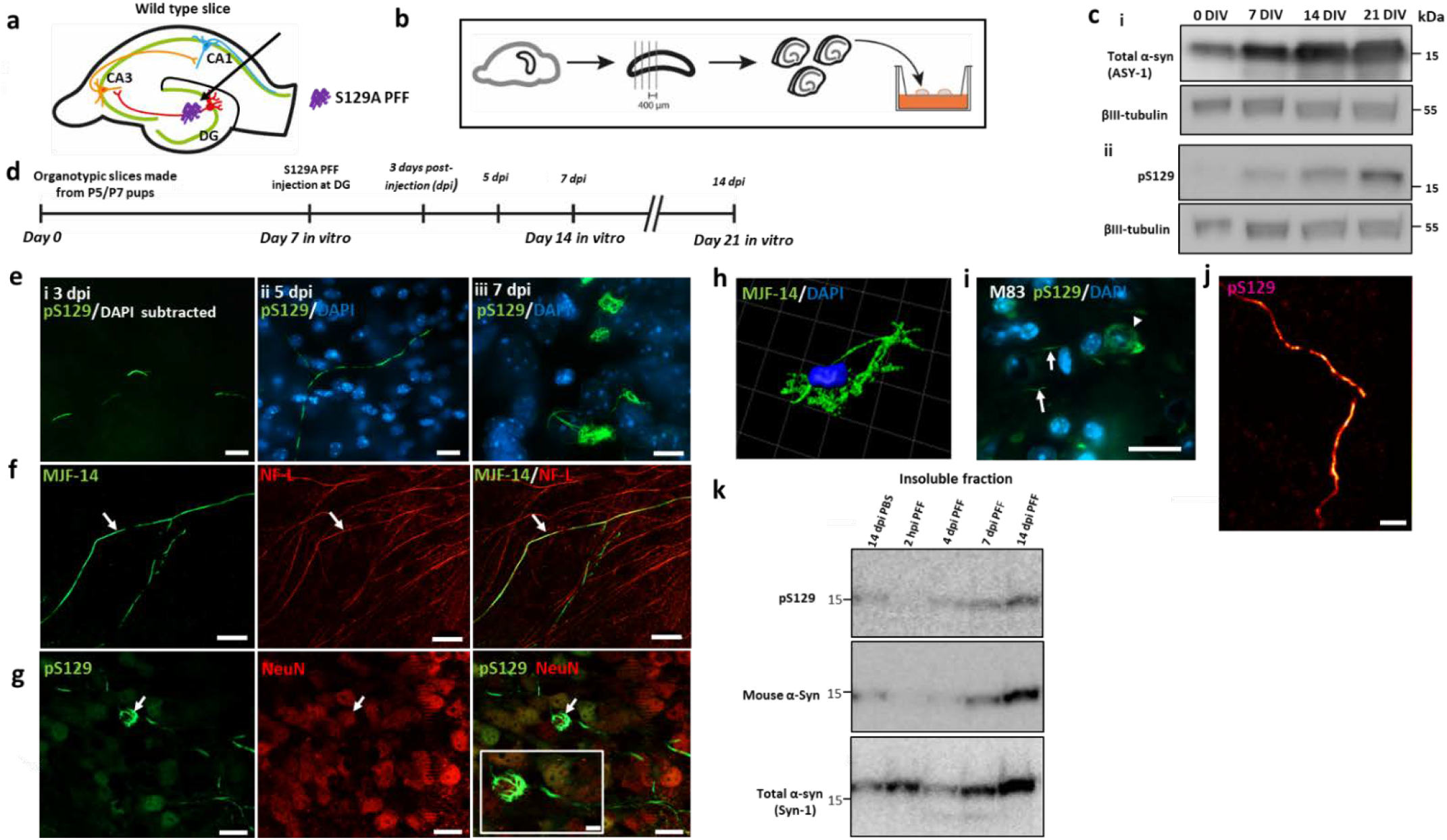
Organotypic mouse hippocampal slice cultures as a model to study seeded α-syn aggregation in the region between DG and CA3. **a**, Bi-synaptic connections of granule cells in DG, where S129APFF were injected to pyramidal neurons in CA3 that subsequently connect to CA1 region. **b**, Organotypic hippocampal slice cultures (OHSC) from mouse pups were cultivated on an air-membrane interface. **c**, Progressive accumulation of total and pS129-α-syn in cultures from wild type mouse pups after 0, 7, 14, and 21 days (DIV) analyzed by immunoblotting. **d**, Experimental flow. **e**, pS129-positive α-syn structures at DG **i**, first recognizable at 3 dpi as short serpentine aggregates that **ii**, coalesce into longer aggregate and **iii**, 7 dpi occurs as fibrillar aggregates around neuronal nuclei, scale bars=20µm. **f**, MJF-14 positive serpentine aggregates colocalize with the axonal neurofilament light chain (NF-L) marker, scale bar= 20µm. **g**, The pS129-positive cell body inclusions are in NeuN-positive neurons, scale bar=20µm. Insert, scale bar=5 µm. **h**, Thread-like cell body inclusion detected by α-syn aggregate-specific antibody MJF14 and reconstructed in 3D by IMARIS software. **i**, The cell body pS129 α-syn pathology in the hindbrain of end-stage h-A53T-α-syn transgenic mice (M83) resembles inclusions in the slice model (panels e,iii & g). **j**, dSTORM image reconstruction of pS129-positive axonal processes within the brain slice. Scale bar=1µm. **k**, Progressive accumulation of insoluble pS129-positive mouse α-syn in PFF-injected slices. Images are representative of more than 3 experiments.

Because endogenous expression of α-syn is a prerequisite for templating α-syn aggregate and subsequent interneuronal spreading, we first determined the development of α-syn expression in the wild type brain slices through a 21-day culture period. The cultivated slices exhibited progressive increase of total α-syn mirroring the postnatal expression in mice (Jakowec, Donaldson, Barba, & Petzinger, 2001) (Fig. 1, c). The phosphorylated pS129 form became detectable after 7 days, after which its level also increased (Fig. 1, c). Based on these data, we chose to initiate the process of aggregation by PFF microinjection in the slice cultured for 7 days *in vitro* (DIV).

As pathological α-syn aggregates are heavily phosphorylated on S129, we used PFFs composed of S129A mutant α-syn that cannot be phosphorylated at this site. This allowed unambiguous detection of endogenous aggregates by pS129-specific antibodies that do not bind the injected S129A seeds. The fate of the exogenous S129A-PFF injected into the brain slices was investigated using tissue from α-syn KO pups allowing us to focus on the injected material. The tissue was analyzed 2 hours, 3 days, and 7 days post injection. To facilitate quantitative immunoblotting, PFFs injected into slices were depolymerized in 7M urea/ 4% SDS loading buffer, which allowed their quantification as a monomeric band. Using the MJFR1 antibody that binds to a C-terminal epitope (118-123),(Schmid, Fauvet, Moniatte, & Lashuel, 2013), we observed a 30% reduction after 3 days with no detectable protein remaining after 7 days. However, using Syn-1 (BD monoclonal antibody) that detects an epitope corresponding to amino acids 91 to 99 (Schmid et al., 2013), we found that the injected PFFs remained in the tissue but as a C-terminally truncated species (Supplementary fig. 1, b). The Syn-1 antibody did not work well for IHC analysis of the PFF-injected α-syn KO slices to localize the truncated species at 5 and 7 days post injection. Staining with MJFR1 revealed that the injected material was confined to a small area close to the injection site in the DG regions 2 hours post injection that disappeared after 7 days as expected from the C-terminal truncation (Supplementary figs. 1, c).

### Injection of S129A-PFF template α-syn aggregation in organotypic hippocampal slice culture

When S129A-PFFs were microinjected into the tissue slice from wild type (C57BL/6) OHSC at DG, pS129-positive aggregates of mouse α-syn were detectable 3 days post injection (dpi). It started as short, serpentine-like inclusions with a diameter of about 0.06 µm (Figs. 1, e,i). The pS129 positive structures were also positive for the aggregate-specific MJF-14 antibody. By 5 dpi, these structures appeared to coalesce into longer serpentine structures, while at 7 dpi, fibrillar aggregates were detected in the cell body around the nucleus (Fig. 1,e,ii,iii). The aggregates located in the cell bodies resembled those observed by IHC in the hypothalamic region of an end-stage hA53T α-syn transgenic mice (Fig. 1,i).

The pS129-positive aggregates in slices were located in axons and neuronal cell bodies as evidenced by their colocalization with the subcellular markers neurofilament light chain and NeuN (Fig. 1,f,g). Upon extraction and western blotting of PFF-injected slices, it was evident that pS129-positive insoluble α-syn species became detectable after 7 days and increased at 14 days. The results demonstrate that endogenous α-syn is converted into insoluble species phosphorylated at Ser129 upon seeding with PFF (Fig. 1,k). Superresolution microscopy revealed the axonal pS129-positive inclusions consisted of smaller structures of more intense immunoreactivity suggestive of discrete α-syn inclusions filling or being moved within the axon (Fig. 1,j).

Injection of monomeric α-syn into WT slices or injection of S129A-PFFs into slices made from α-syn KO pups induced no aggregation at 7 dpi (Fig. 2,a,i,ii). This shows that the aggregation process depends on preformed fibrillar seeds and presence of endogenous α-syn.

**Fig. 2:**
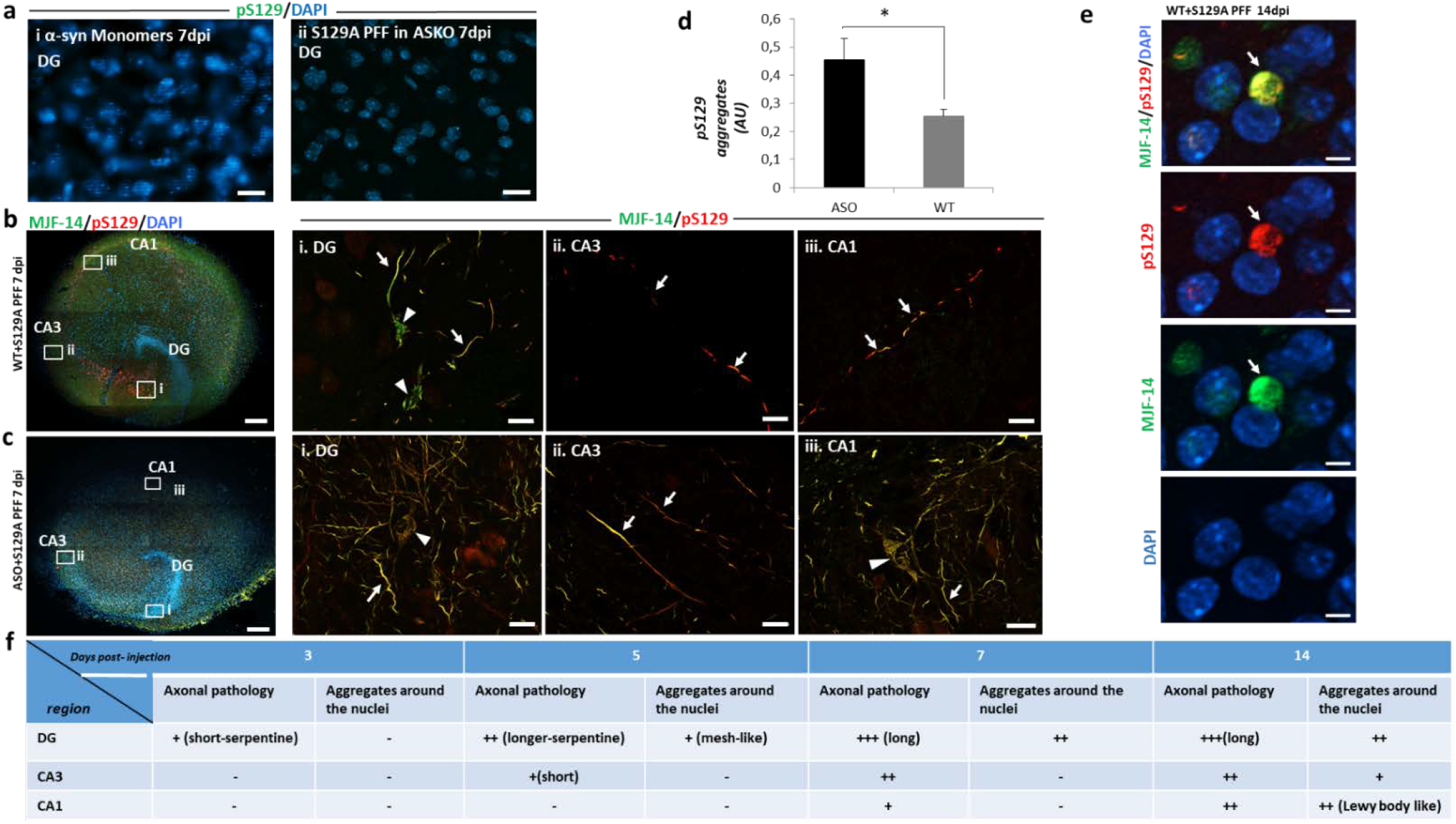
Trans-synaptic spreading of α-syn aggregate-pathology from DG via CA3 to the CA1 region of the hippocampus depends on expression levels of α-syn. **a**, No aggregation is induced by injecting i. Monomeric α-syn in wt slices or ii. S129A-PFF in slices from α-syn knockout(ASKO) pups, scale bar=20µm. **b**, Composite images of immunostaining with aggregated (MJF-14, green) and pS129-α-syn(red) 7 dpi from wt OHSC, scale bar=200µm. Areas from DG, CA3, and CA1 regions indicated are magnified in panels i, ii, and iii, scale bar=20 µm. Axonal aggregates (arrows) are present in all three regions, while cell body inclusions (arrow heads) are present only in DG by 7dpi. **c**, Composite image of immunostaining with MJF-14 and pS129 aggregates 7dpi in OHSC from h-α-syn transgenic mice (ASO), scale bar=200 µm i, ii, Florid pS129-α-syn and iii, faster progression with cell body inclusions in the CA1 region, scale bar=20 µm. **d**, Quantification of pS129-α-syn aggregate fluorescence signals in total slices from PFF-injected WT and ASO slices. Bars represent mean ± SD, n=3. Unpaired Student’s T-test, p-value=0.019. **e**, Immunostaining with pS129 and MJF-14 at CA1 region of wild type slices 14dpi of PFFs. Showing more compacted spherical cytoplasmic inclusions, resembling Lewy body. Scale bar= 5 µm. **f**, Schematic demonstration of progressive evolution from short into longer serpentine axonal inclusions in DG regions that spread to CA3 and CA1 regions with cell body inclusions appearing at later stages when axonal pathology is established in the region. Images are representative of more than 3 experiments.

### Application 1: Demonstrating S129A-PFF templated α-syn aggregation spreads by interneuronal processes from the DG region via CA3 to CA1 region

Seeded aggregation of endogenous α-syn in the DG regions was observed after 3 dpi, but the first interneuronal spreading from DG to the CA3 and CA1 regions appeared as axonal inclusions after 5 to 7 days (Fig. 2,b). Neuronal cell body inclusions became detectable after 2 weeks in CA1 region (Supplementary Fig. 2), that in comparison to the aggregates found in the cell bodies of DG, were more compacted as spherical cytoplasmic inclusion with immunoreactivity to both pS129 and MJF-14 antibodies, suggesting the development of Lewy body-like inclusion at this stage in the culture (Fig. 2, e & Supplementary fig.3). Both seeding and spreading developed more quickly and became more prominent when the endogenous α-syn level was increased, as demonstrated in slices from mThy-1-human-α-syn transgenic pups, where cell body inclusions were detected at the CA1 region by 7dpi (Fig. 2,c,d). Microinjection of PFFs was essential for the ordered interneuronal spread of PFF-templated aggregation from DG to CA3 and CA1, because application of PFF solution to the surface of the slice resulted in development of pS129-positive structures predominantly in the periphery of the slice (Supplementary fig. 4). This pattern is most likely caused by fluid flow across the slice surface.

Having established that seeded α-syn aggregate pathology can spread anterograde from the DG to the CA1 region in the hippocampal slice model, we wanted to determine if retrograde spreading from the CA1 to the DG region also could occur. When slices were injected with S129A-PFFs in the CA1 region, pS129-positive aggregates were observed at CA1 and in the area between CA1 and CA3 after 10 days, but no aggregates were found at DG suggesting retrograde spreading is less efficient compared to the anterograde spreading (Fig. 3,a,i,ii,iii).

**Fig. 3:**
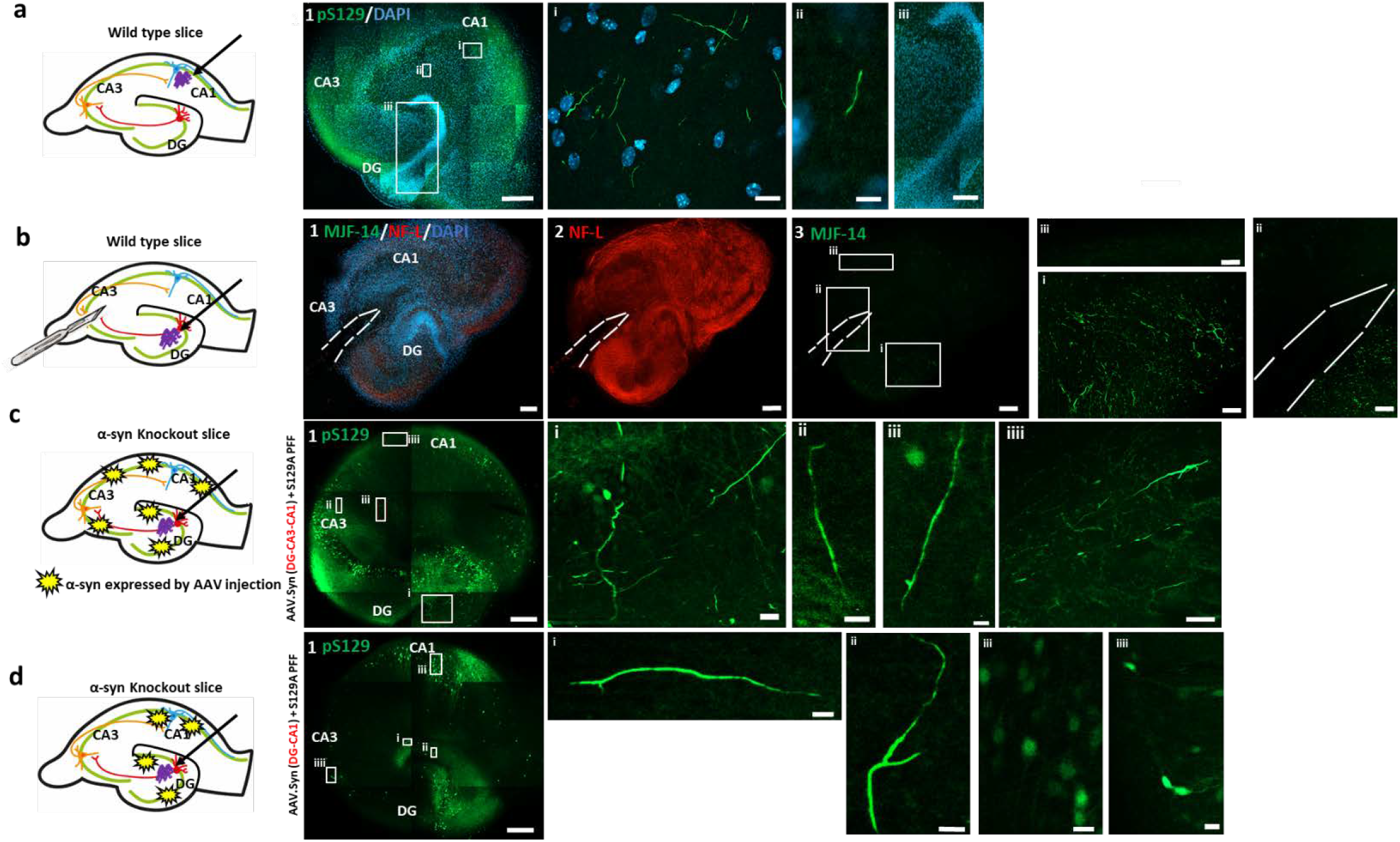
Application I. Demonstrating anterograde trans-synaptic spreading is the predominant route for progression of α-syn-aggregate pathology spreading from DG via CA3 to the CA1 region of the hippocampus using surgical and viral transgene methods. **a**, Schematics of PFF injection in CA1 in WT OHSC. 1, Composite image 10 dpi of S129A-PFFs at CA1, scale bar=200 µm. i: Inset from CA1 showing pS129-positive aggregates, scale bar=20 µm. ii: Inset from region between CA1 and CA3, scale bar=5 µm. iii: Inset from DG with no aggregates, scale bar=50 µm. **b**, Schematics showing transection of axonal projections between DG and CA3 that block spreading of α-syn aggregate pathology from DG to CA1. The surgical destruction of the tissue is demonstrated by absent **1**, nuclei and **2**, axonal marker NF-L, and **3**, MJF-14 staining. 1,2,3 scale bar=200 µm. i, magnified image showing aggregates at DG. ii, aggregates found distal to the cut but not proximal to it. iii, No aggregates found at CA1 region. i,ii,iii, scale bar=50 µm. **c**, Schema showing how wt**-**α-syn was expressed in ASKO slices by AAV vectors injected in DG, CA3, and CA1. **1**, α-syn expression in DG, CA3, and CA1 supports spreading of aggregated pS129 pathology to CA1 upon PFF injection in DG. Scale bar in panels: 2=200µm, i=20 µm, ii&iii=10 µm. Note the strong AAV-dependent expression of pS129 in some neuronal nuclei. **d**, Schema showing that wt**-**α-syn was expressed in ASKO slices in DG and CA1 only. **1**,Absent α-syn expression in the CA3 hindered the aggregates spreading to CA1 at 7dpi. Scale bar=200 µm. **i,ii**,pS129-positive aggregates are detectable at DG. **iii**,no pS129-positive aggregates found at CA1 region. **iv**,few neurons show nuclear expression of pS129 at CA3. Scale bar= 20 µm i, ii, iii=10 µm. Images are representative of more than 3 experiments.

To test the hypothesis that the identified interneuronal spreading is trans-synaptic in the slice model, a surgical cut was made in the region between DG and CA3 through the whole thickness of the slice immediately after PFF injection. This would block the axonal transport required for spreading through the synaptically connected DG and CA3 regions. Loss of axonal connectivity between DG and CA3 in the slice blocked the formation of the MJF14-and pS129-positive aggregates distal to the lesion compared with control slices with an intact connection through the DG-CA3-CA1 regions of the slice (Fig. 3,b).

True trans-synaptic spreading of templated α-syn aggregate pathology requires expression of endogenous α-syn in all the connected neurons as α-syn is required to amplify the small amount of seeds taken up by receiving neurons to facilitate further release of new seeds to the next neuron in the pathway. To investigate this hypothesis, we used α-syn KO slices that do not allow development of pathology upon PFF injection (Fig. 2, a,i). In these slices, human α-syn was expressed in neurons by AAV vector driven by the synapsin1 promoter at all DG, CA3 and CA1 regions (Fig. 3, c). Successful expression was confirmed with immunostaining of AAV injected slices using total α-syn (MJFR1) and pS129 antibodies. α-syn and pS129 expressing neurons were detected at the targeted regions with pS129 positive diffuse nuclei and cell bodies whereas MJFR1 staining revealed α-syn was efficiently sorted to punctate-structures reminiscent of nerve terminals (Supplementary Fig. 5, a, i & ii). pS129 antibody were used for detection of 129A PFF induced aggregates. These more pale signals are clearly distinct from the brighter inclusions that evolved upon injection of PFF (Fig. 3, c,i,ii,iii).

Injection of α-syn-expressing vectors in the DG, CA3, and CA1 regions of the KO slice allowed interneuronal spreading of α-syn aggregate pathology from the PFF injection site in the DG to the synaptically connected CA3 and CA1 regions (Fig. 3, c,i,ii,iii) as observed in wild type hippocampal slices (Fig. 2, b). The pathology progressed further by 14 dpi, with the perinuclear pS129-positive aggregates found in the CA1 region (Supplementary fig. 5, b), as discussed for wild type slices injected with PFFs. To abrogate the hypothesized sequential trans-synaptic spreading from neurons in DG to CA3 and then CA1, we only injected the human α-syn-expressing AAV vector into the DG and CA1 regions but not in the CA3 neurons, to leave CA3 region without any α-syn expression (Fig. 3, d). No spreading to the CA1 region was detectable by 7 dpi upon injection of S129A PFF into the DG region, although a prolific pS129-positive aggregation developed in the DG region (Fig. 3, d, i, ii, iii). Even at 14dpi, no pathology was detectable in the CA1 region, showing only the diffuse nuclear staining (Supplementary Fig. 5, c). This supports the hypothesis of sequential trans-synaptic spreading of α-syn aggregate pathology.

### Application 2: Using organotypic hippocampal slices from α-syn KO mice and AAV vectors to demonstrate α-syn aggregation and spreading occur independently of its phosphorylation on Ser-129

Phosphorylation of serine-129 in α-syn represents a hallmark for pathological α-syn inclusions in the human brain. However, its pathophysiological role is contested with respect to its role in aggregation and cytotoxicity(Gorbatyuk et al., 2008), (McFarland et al., 2009),(Chen & Feany, 2005). To investigate the role of this post-translational modification in the trans-synaptic spreading of seeded α-syn aggregation using the new model, we expressed a non-phosphorylatable S129G mutant and wt α-syn using AAV vectors in α-syn KO slices, and compared the spreading of MJF-14 positive aggregates for both variants (Fig. 4, a). S129G and wt α-syn were expressed at comparable levels in the KO slices as determined by immunoblotting, but only the wt protein was as expected phosphorylated on S129 (Fig. 4, b). Both transgenes supported the development of MJF-14 positive aggregates in the DG regions that spread to the CA1 region when S129A-PFFs were injected into the DG (Fig. 4, c, d). However, only the wt form of α-syn was phosphorylated at S129 (Fig. 4, e, f). The S129 phosphorylated and non-phosphorylable aggregates appeared comparable with respect to axonal and cell body localization. It can thus be concluded that phosphorylation of S129 in α-syn is not critical for interneuronal spreading of the templated aggregates pathology. The organotypic hippocampal slice model is thus a versatile tool that allows investigation of the role of site-specific post-translational modification in spreading of seeded α-syn in a time and cost-scale that is much smaller than if one had to develop transgenic animal models on an α-syn KO background.

**Fig. 4:**
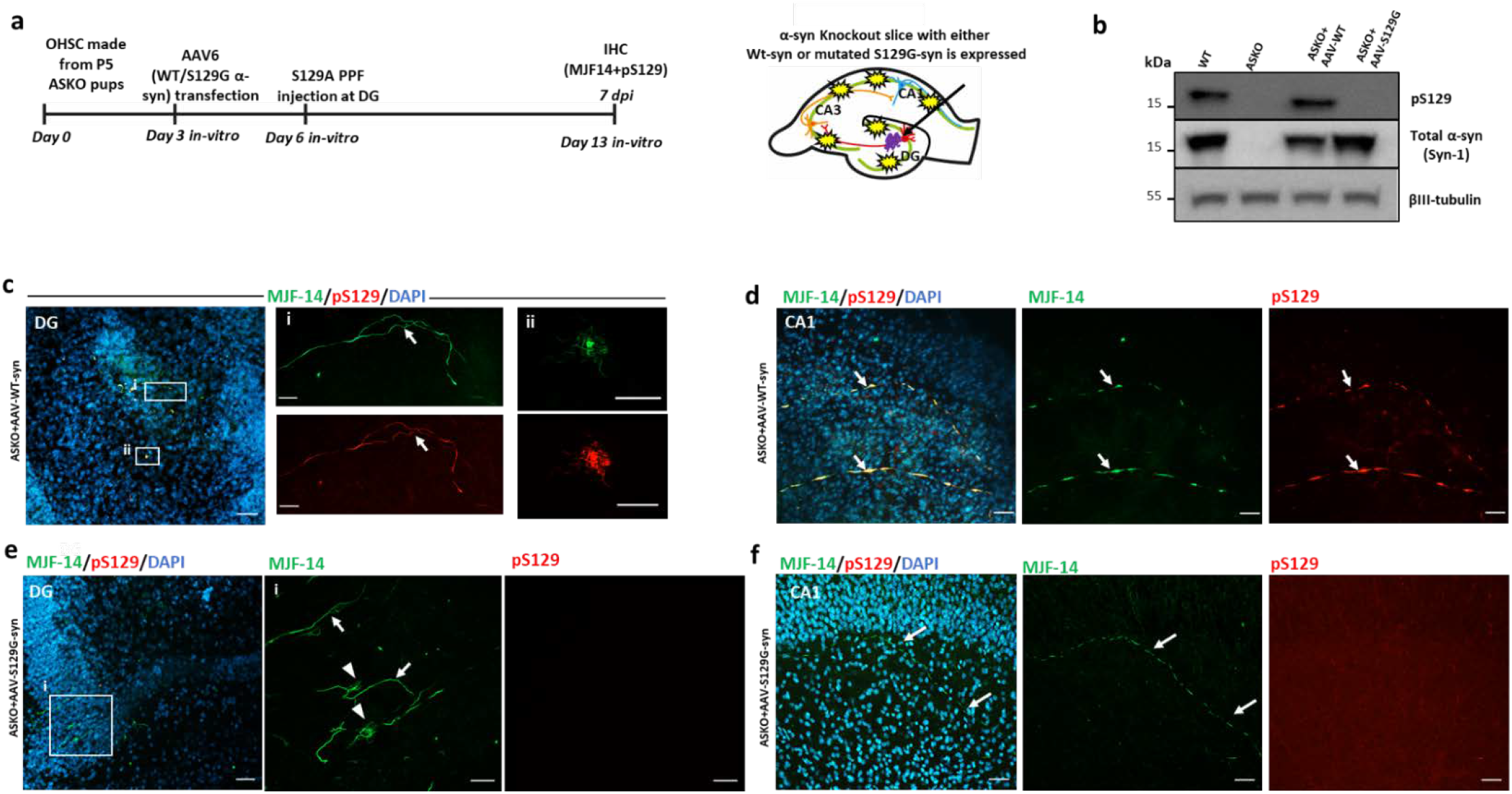
Application II. Demonstrating phosphorylation of S129 on α-syn is not a prerequisite for seeding α-syn aggregation and trans-synaptic spreading in hippocampal slices. **a**, Schema of experiment where neuronal expression of either wt or non-phosphorylatable S129G-α-syn is established prior to initiation of templated α-syn aggregation by injection of S129A-PFF. **b**, Validation of virally mediated WT- and non-phosphorylatable S129G-α-syn expression in slices from α-syn knockout mice (ASKO) using antibodies against total and pS129 α-syn. **c**, Expression of wt α-syn supports establishment of pS129-positive aggregate pathology, scale bar=50 µm in i, axonal aggregates, scale bar=20 µm and ii, cell bodies inclusions at DG, scale bar=5 µm. **d**, The pathology spreads to the CA1 region scale bar=50 µm. **e**, Despite not being phosphorylated on S129, expression of S129G-α-syn still supports establishment of MJF-14-positive aggregate pathology in the DG, scale bar=50µm that **i**, is present in axons (arrow) and cell bodies (arrow head), scale bar=20 µm. **f**, This nonphosphorylated aggregate α-syn pathology spreads to the CA1 region, scale bar= 50 µm. Images are representative of more than 5 experiments.

## 4. Discussion

The correlation of PD symptomatology with the progression of the pathological α-syn aggregate-containing Lewy bodies formed the basis for the Braak hypothesis (Braak et al., 2003). This hypothesis was corroborated by the demonstration of Lewy body pathology in fetal neurons transplanted into the striatum of PD patients 11-16 years prior to their death (Brundin, Melki, & Kopito, 2010). Since then, the general hypothesis of a prion-like spreading of templated misfolded proteins that comprised diseases like Creutzfeldt-Jakob disease and Alzheimer’s disease was expanded to include other disease as PD, dementia with Lewy bodies (DLB), and multiple systems atrophy (MSA), where seeds of aggregated α-syn are considered the culprit (Jucker & Walker, 2018),(Aguzzi, Baumann, & Bremer, 2008),(Prusiner et al., 2015). *In vivo* models of seeded α-syn aggregation are slow and costly, and *in vitro* cell-based models developed to facilitate mechanistic investigation and drug screening lack the complexity of nervous tissue (Gribaudo et al., 2019),(Volpicelli-Daley, Luk, & Lee, 2014), (Smits et al., 2019).

In this article, we present a novel *ex vivo* brain tissue culture model that can serve as a translational step between cell-based and *in vivo* models. The model takes advantage of the organotypic hippocampal slice culture that for decades has been used for electrophysiological analyses(Dunwiddie & Lynch, 1978). It possesses three synaptically connected neuronal populations located in a developmentally evolved brain structure embedded in an active matrix of glia cells, and it is thus outperforms currently available two- and even three-dimensional *in vitro* culture systems (Smits et al., 2019),(Amin & Paşca, 2018). The model is based on microinjection of α-syn PFFs into the DG. PFFs seed a progressive, templated aggregation of endogenous mouse α-syn that spreads through CA3 to CA1 neurons. Application of the PFFs by microinjection is critical for the model because simple application of PFFs on top of the slice only induced development of aggregate-containing neurons in the periphery of the slice and not the interconnected neurons of the DG, CA3, and CA1 circuit.

The use of S129A-PFF, which cannot be phosphorylated at S129, and detection of cellular aggregates by pS129-α-syn antibody allowed the unequivocal conclusion that inclusions formed in the tissue were made *de novo* and do not merely represent the injected material. These inclusions shared the disease-associated epitopes with aggregates present in human brain affected by synucleinopathies as demonstrated by the binding of pS129-α-syn antibodies and the conformation-specific α-syn aggregate antibody MJF-14 (Sampson et al., 2018). The PFF induced inclusions had demonstrated different patterns that were topology- and time-dependent (Supplementary fig. 3). Interestingly, the aggregates observed at DG by 7 and 14 dpi resembled filamentous structures that wrapped the nucleus. However, more compacted, often spherical, cytoplasmic inclusions developed in CA1 region at 14dpi, suggesting Lewy body-like inclusions can development in this region. This concurs with findings demonstrating Math2 expressing neurons in the CA1 were more affected by α-syn pathology following seeding with PFFs in vivo, in contrast to DG neurons, which are relatively spared (Luna et al., 2018).

No pathology developed when PFFs were injected in α-syn KO slices. This demonstrates that neuronal α-syn expression is essential for the templating of α-syn aggregate pathology in the model in accordance with previous *in vivo* models (Luk et al., 2012).

In agreement with the previous finding that the development and progression of α-syn pathology in humans depends on the level of α-syn expression, where faster disease progression in patients with duplications and triplications of the α-syn-encoding SNCA gene was reported (Konno, Ross, Puschmann, Dickson, & Wszolek, 2016),(Singleton et al., 2003), the development and progression of α-syn in the hippocampal slice model were enhanced in slices from α-syn-transgenic pups compared with wild type pups..

The directionality of the pathology spreading remains unclear because of the complex spreading pattern observed in the brain after inoculation of seeds in the striatum, hippocampus or olfactory bulb (Rey et al., 2018),(Masuda-Suzukake et al., 2013). However, a component of retrograde axonal transport is likely to transfer seeds from the gut to the vagal nucleus, as demonstrated in rodents (Holmqvist et al., 2014). We used the model to answer the question of whether there is a preferred direction of spreading of templated aggregation in hippocampal tissue by comparing the spreading upon injection of PFF into the DG or CA1. Anterograde transfer from the DG to the CA1 region was the preferred mode, as demonstrated by the absence of spreading from the CA1 region to the DG after 10 days despite florid local templating of α-syn to an aggregated form at the CA1 region. The hypothesis of anterograde trans-synaptic spreading was corroborated by the fact that spreading was blocked by cutting the axons in the CA3 region. To demonstrate spreading not only requires an intact synaptic connection but also a sufficient level of α-syn expression to allow a seeding dependent aggregate formation. We therefore used hippocampal slices from α-syn KO pups in which neuronal α-syn expression was established locally using AAV vector. The α-syn KO slices were only able to support templated spreading from DG to CA1 when human α-syn expression was established in all connected regions, DG, CA3, and CA1. Inversely, no spreading occurred if the slices lacked α-syn expression at the CA3 region. This experiment also demonstrates that templated aggregation in the CA1 region is due to a transfer of templated pathological α-syn aggregates through the circuit, which requires renewed templating in each recipient population and not just a transfer of injected PFF through the circuit

The viral vector approach to the expression and silencing of genes of interest makes our slice model a versatile tool for constructing tailor-made slices where candidate genes are modulated. As proof of concept, we asked if phosphorylation on S129 is necessary for seeded aggregation and spreading in brain tissue because this post-translational modification represents a biomarker of aberrant α-syn aggregates in cells and tissue. To answer this question, we used AAV vector to reconstitute expression of WT and non-phosphorylatable S129G mutant human α-syn in α-syn KO slices and injected them with PFFs in the DG region. The successful production of a slice solely expressing S129G α-syn allowed us to conclude that phosphorylation at S129 was not a prerequisite for initiation of α-syn aggregation or its interneuronal spreading. The approach of transgenic expression of α-syn species in KO tissue can be extended to investigate the role of other post-translational α-syn modifications like truncations, ubiquitinations on specific lysines, or N-terminal acetylation; and it can be used for validating proteins involved in spreading process.

The model has some limitations; firstly, anatomic variations among individually cultivated slices sectioned through the small hippocampal structure can pose a challenge. However, random selection and increasing the number of slices used per experiment can overcome the influence of this variation in the final results. Secondly, the sparse induction of aggregates by PFFs in only certain neurons when wild type slices are used makes it more difficult to study the conspicuous toxicity of aggregates in slices densely packed with neurons but may also open for studying the effect of different α-syn aggregate strains on uptake, seeding, and spreading. It may also make it possible to address factors governing selective neuronal vulnerability and thus resemble *in vivo* conditions in the sick brain, where only vulnerable neurons rather than the whole population show synuclein pathology.

## 4. Conclusion

This study presents a novel *ex vivo* brain tissue model for studying seeded α-syn aggregation and interneuronal spreading in circuitry-connected neurons. This model is superior to previous *in vitro* models with regard to replicating the hypothesized pathophysiologic neuronal handling of α-syn aggregates as only the first recipient neurons in DG are challenged by *in vitro* formed aggregates. The further spreading of seeding competent species represents novel *in cellulo* generated aggregates as demonstrated by the absence of aggregates in slices from synuclein α-syn KO slices and the absence of spreading from DG to CA1 when no α-syn was expressed in the CA3 neurons. These underlying mechanisms may thus be operating in diseases like PD, Lewy body dementia, and Alzheimer’s disease, which exhibits significant Lewy body accumulation. The speed by which the spreading occurs combined with the ability to manipulate the model genetically by using hippocampal tissue of pups from transgenic mouse lines and AAV viral vector allows for testing a host of pro- and anti-degenerative genes including risk genes identified by GWAS studies. With respect to posttranslational modifications, we established that phosphorylation at S129 not is a prerequisite for aggregation or spreading in mouse tissue. It is thus an innovative alternative to transgenic animals, being significantly easier, faster, and less costly; most importantly, it is well in line with the “3R” concept values, replacement, reduction and refinement, regarding use of animals in experiments. It also provides opportunities for novel, attractive, and beneficial applications based on using different genetic mouse lines, viral vector-based gene regulation, super-resolution microscopy, live imaging and electrophysiological recording, and pharmacological treatment.

## Acknowledgement

The authors would like to thank Simon Mølgaard Jensen, PhD, Department of Biomedicine, AU for his help with 3D image reconstruction using IMARIS program and Björn Anzelius, Research engineer at Brain Repair and Imaging in Neural Systems (BRAINS), Lund University, Sweden.

We would also acknowledge the following funding sources: The Lundbeck Foundation grants R248-2016-2518 for Danish Research Institute of Translational Neuroscience - DANDRITE, Nordic-EMBL Partnership for Molecular Medicine, and R223-2015-4222. Michael J Fox Foundation grant 12028.01. Cultural Affairs and Mission Sector, Ministry of Higher Education, Arab Republic of Egypt, The European Research Council ERC CoG-724489 and The Swiss National Science Foundation (Sinergia) CRSII3_154461.

